# Observation of a single protein by ultrafast X-ray diffraction

**DOI:** 10.1101/2022.03.09.483477

**Authors:** Tomas Ekeberg, Dameli Assalauova, Johan Bielecki, Rebecca Boll, Benedikt J. Daurer, Lutz A. Eichacker, Linda E. Franken, Davide E. Galli, Luca Gelisio, Lars Gumprecht, Laura H. Gunn, Janos Hajdu, Robert Hartmann, Dirk Hasse, Alexandr Ignatenko, Jayanath Koliyadu, Olena Kulyk, Ruslan Kurta, Markus Kuster, Wolfgang Lugmayr, Jannik Lübke, Adrian P. Mancuso, Tommaso Mazza, Carl Nettelblad, Yevheniy Ovcharenko, Daniel E. Rivas, Amit K. Samanta, Philipp Schmidt, Egor Sobolev, Nicusor Timneanu, Sergej Usenko, Daniel Westphal, Tamme Wollweber, Lena Worbs, P. Lourdu Xavier, Hazem Yousef, Kartik Ayyer, Henry N. Chapman, Jonas A. Sellberg, Carolin Seuring, Ivan A. Vartanyants, Jochen Küpper, Michael Meyer, Filipe R.N.C. Maia

## Abstract

The idea of using ultrashort X-ray pulses to obtain images of single proteins frozen in time has fascinated and inspired many. It was one of the arguments for building X-ray free-electron lasers. According to theory^1^, the extremely intense pulses provide sufficient signal to dispense with using crystals as an amplifier, and the ultrashort pulse duration permits capturing the diffraction data before the sample inevitably explodes^2^. This was first demonstrated on biological samples a decade ago on the giant mimivirus^3^. Since then a large collaboration^4^ has been pushing the limit of the smallest sample that can be imaged^5,6^. The ability to capture snapshots on the timescale of atomic vibrations, while keeping the sample at room temperature, may allow probing the entire conformational phase space of macromolecules. Here we show the first observation of an X-ray diffraction pattern from a single protein, that of *Escherichia coli* GroEL which at 14 nm in diameter^7^ is the smallest biological sample ever imaged by X-rays, and demonstrate that the concept of diffraction before destruction extends to single proteins. From the pattern, it is possible to determine the approximate orientation of the protein. Our experiment demonstrates the feasibility of ultrafast imaging of single proteins, opening the way to single-molecule time-resolved studies on the femtosecond timescale.

---

X-ray free-electron lasers (XFEL) have transformed the study of ultrafast phenomena at the atomic level, from transient room-temperature superconductivity^8^ to the fastest processes following water ionisation^9^. This has also been the case in structural biology with the birth of serial femtosecond crystallography (SFX)^10^ and more recently the development of time-resolved SFX^11^. Yet the requirement of crystals is limiting as demonstrated by the spectacular development in cryo-electron microscopy (cryo-EM)^12^. More importantly, the need to synchronise all unit cells in a crystal makes photo-activation the only feasible trigger for ultrashort timescales. It also prevents the observation of individual molecular behaviour, e.g. multiple conformations. Currently, cryo-EM is the method of choice for high-resolution single-molecule time-resolved studies, but it is limited to millisecond timescales due to the time it takes to freeze the sample and collect the data^13^. By bypassing these limitations, femtosecond X-ray diffractive imaging (FXI)^1^ has the potential to observe single-molecules with sub-picosecond time-resolution and, due to the higher sample temperature, may allow sampling from a broader conformational landscape.

The chaperonin GroEL is an abundant molecular chaperone and, together with its cofactor GroES, is important in the folding of a large range of proteins^14^. *E. coli* GroEL is a 14-mer formed by two heptameric subunit rings, totalling ~800 kDa and arguably the most studied chaperonin. It was also one of the first large macromolecular complexes to be successfully measured by native mass spectrometry^15^ and is nowadays often used as a benchmark to demonstrate the resolution of new systems^16–18^. Its size and availability also made it an early target for single-particle cryo-EM studies^19,20^. These characteristics along with the extensive body of available knowledge and distinctive shape, recognizable even at low resolution, make GroEL an ideal prototype system for single-particle X-ray diffraction.

Despite continuous progress in FXI, no single-protein diffraction has ever been measured, and studies have been limited to more strongly diffracting samples, such as viruses^21^ and cells^22^. In this paper we present the first interpretable X-ray diffraction signal from a protein complex and with it demonstrate the principle of diffraction before destruction^2^ at the protein scale. This opens the doors to ultrafast studies on single protein molecules making use of the extraordinary brightness and time-resolution of XFELs.

The experiment was performed at the Small Quantum Systems (SQS) scientific instrument of the European XFEL (EuXFEL) facility in Schenefeld, Germany^23^. GroEL particles were exposed to femtosecond soft X-ray pulses from the EuXFEL at a photon energy of 1200 eV and average pulse energy of 6.5 mJ.

Individual GroEL particles, characterised by a differential mobility analyzer (DMA) (**Fig. S1**) and cryo-EM (**Figs. S2-5**), were transferred from solution to the gas phase using an electrospray setup^24^ in which a charged jet of the sample in liquid generated droplets of around 110 nm in diameter in the presence of an inert gas mixture of CO_2_ and N_2_ surrounding the jet (**Fig. 1**). These droplets were then neutralised and focused through an aerodynamic lens^25^ creating a thin stream of particles. Most or all of the volatile buffer solution evaporated during the process and a stream of mostly dry particles reached the interaction region.

**Fig. 1.**
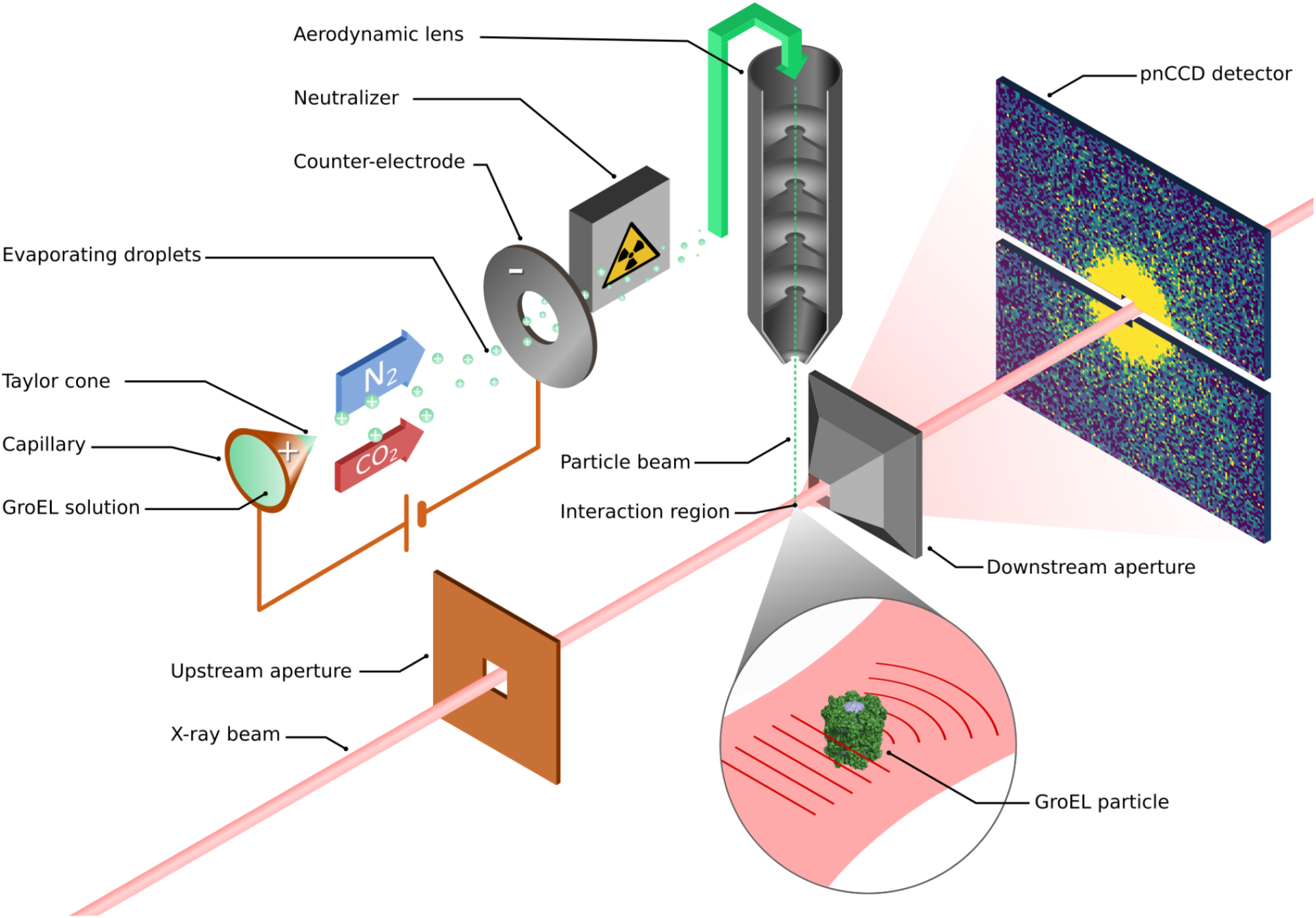
Experimental setup. A solution containing GroEL particles, each roughly a cylinder 14 nm in diameter and height, is aerosolized, using electrospray ionisation followed by neutralisation, and focused into a thin stream using an aerodynamic lens. The stream is then intersected with the path of the XFEL beam and the diffracted signal is collected on a pair of pnCCD detectors downstream of the interaction region. To minimise the amount of background, the beam is cleaned up by apertures both before and after the interaction region^26^.

Diffraction data were collected with a pnCCD detector consisting of two detection planes^27^ placed 150 mm downstream of the interaction region (**Fig. 1**). The resolution limit of this setup is 4 nm due to the detector’s numerical aperture. Only a small fraction of the X-ray pulses will intersect with one of the injected particles in what is called a hit. The majority of the detector readouts therefore only contain background, which arises mainly from the injection gas but also from the beamline itself.

The gas used in the electrospray injection setup created two types of experimental background: fluorescence and elastically scattered photons. The fluorescence has a photon energy of 277 eV, 392 eV and 525 eV respectively from the carbon, nitrogen and oxygen K_α1_-shell, compared to the incoming photons of 1200 eV. The energy resolution of the pnCCD detector of 40 eV^28^ allows us to discriminate between the fluorescence and elastic scattering for all pixels that receive at most one photon (**Fig. S6**), a condition that was generally fulfilled in this experiment.

In contrast to the fluorescence background, it was not possible to filter out the elastic scattering from the gas since it has the same photon energy as the signal. The same is also true for the so-called beamline background – photons resulting from the interaction of the X-rays with elements of the beamline. To quantify the different sources of background we collected data both with the injection off and the injection turned on but without a supply of sample. This showed that the injection gas contributed on average 17,600 photons per diffraction pattern, compared to the beamline contribution of only 86 photons per diffraction pattern on average (**Fig. 2**).

**Fig. 2.**
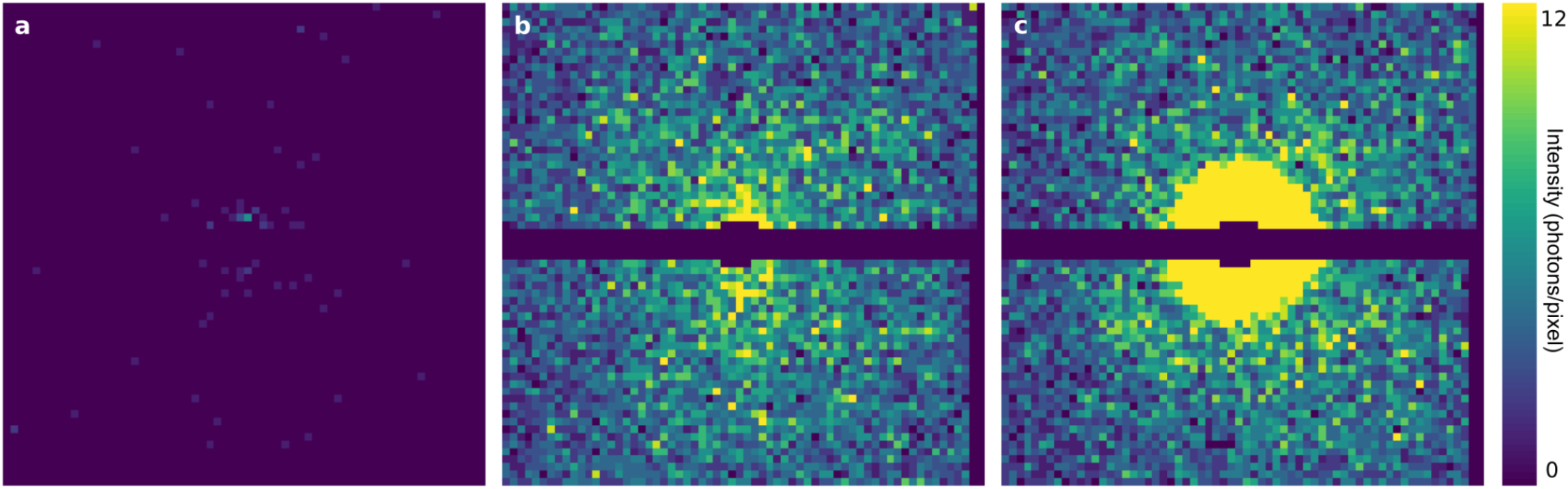
Experimental diffraction data. **a**, Average beamline background, i.e. background with gas from injection turned off, plotted with Poisson noise. **b**, Average measured background, plotted with Poisson noise. **c**, Measured diffraction of a GroEL molecule. All patterns are downsampled to 64×64 to make features more visible on this figure.

The EuXFEL delivers its pulses in 10 pulse trains per second and with a MHz repetition rate within each train^29,23^. Because a detector based on CCD technology is not capable of providing a MHz image read-out rate within the pulse train, we were limited to one readout per train which severely limited the data collection rate. As a consequence, the number of hits was low and only one of them had the combination of high signal-strength and favourable orientation that made it recognizable as GroEL (**Fig. 2c**). Nevertheless, a deviation from the circular symmetry in the first fringe is clear and consistent with the barrel-shaped structure of GroEL. To verify that the pattern originates from a GroEL particle, we compared it with simulated diffraction data from the structure of GroEL determined by X-ray crystallography^30^. This comparison does however have three problems: (1) the orientation of the molecule that gave rise to our pattern is unknown; (2) the centre of the diffraction pattern is uncertain; (3) our diffraction data is a combination of signal and background.

We addressed problems (1) and (2) by applying a template matching scheme where many diffraction patterns were simulated in orientations sampling the full three-dimensional diffraction space with an accuracy of 7 degrees. These patterns were then translated both horizontally and vertically to cover the different possible centre positions. In total, the experimental pattern was compared to 1.2 million simulated and translated patterns.

To handle problem (3) we first summed up the average background from one of the runs where gas but no sample was injected. For each comparison under the template matching, the pattern was fitted to a linear combination of the average background and the template pattern. The best-fitting background-template combination is shown in **Fig. 4a**.

Even this best-fitting background-template combination does not match the experimental pattern very well. The sum of the residual errors between the pattern and the simulation is 254 photons compared to 180 photons which would have been expected if Poisson noise was the only cause for the discrepancy. A hint at an explanation can be found by observing that the first fringe in the simulation is significantly stronger than in the experimental pattern. This indicates that the simulation has too many low-resolution high-contrast elements. This suggests that the hollow centre of the barrel-shaped protein in the simulation is fully or partially filled in the particle that gave rise to the pattern.

We identify three possible origins for this density: (1) It is possible that not all of the water evaporated from the sample during injection, in particular water molecules that are less exposed to the surface. (2) 2D class averages from our cryo-EM measurements (**Fig. S4**) show some density inside the barrel higher than the surrounding water. This density is most likely protein. (3) Depending on the size of the initial solvent droplet there will be a considerable amount of contaminants left on the sample after evaporation. This contributes to the peak at 11 nm observed in DMA data (**Fig. S1**) and could explain the extra density. At the resolution available in this experiment we cannot determine if any of these hypotheses is correct. We can, however, test the theory that extra density within the centre of the protein can explain the observed data.

To do this we created six different density models (**Fig. 3**) by adding varying amounts of water to the hollow centre or the surrounding groves in the protein. We then repeated the template matching with each of them, knowing that similar models filled with broken proteins or salt would give indistinguishable results. Five of the models fill up the hollow core of the protein at varying proportions, which is what our earlier interpretation of the data suggests. As a control, we also include a density model where only the barrel edges are hydrated and the core is empty.

**Fig. 3.**
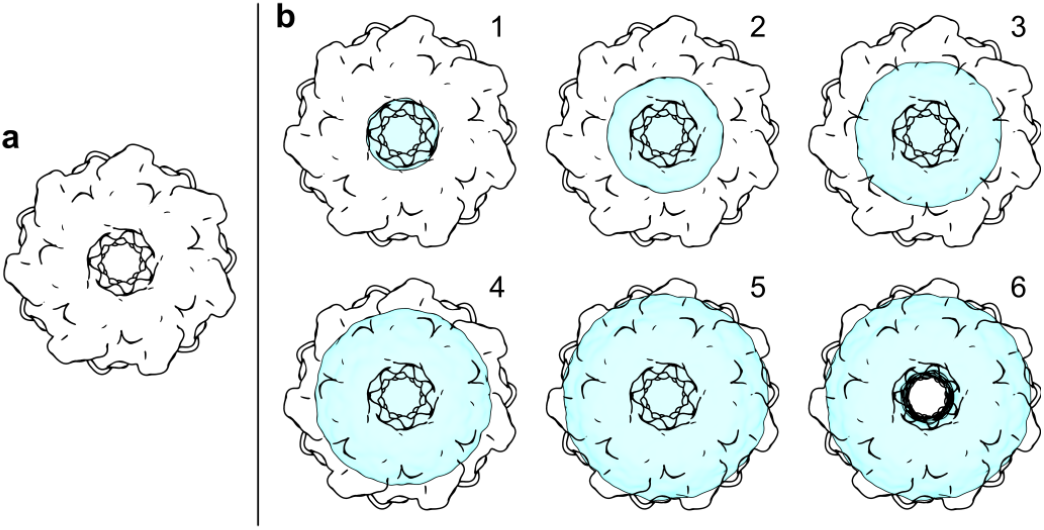
Density models. The original structure **a**, and six models with added density, **b**, were compared to the recorded diffraction intensity. The density is modelled as water. The weight of water, in relation to the weight of the protein, for models 1 to 6 is 13%, 24%, 37%, 51%, 69% and 54% respectively. All models fill the hollow core of GroEL except for model 6.

**Fig. 4.**
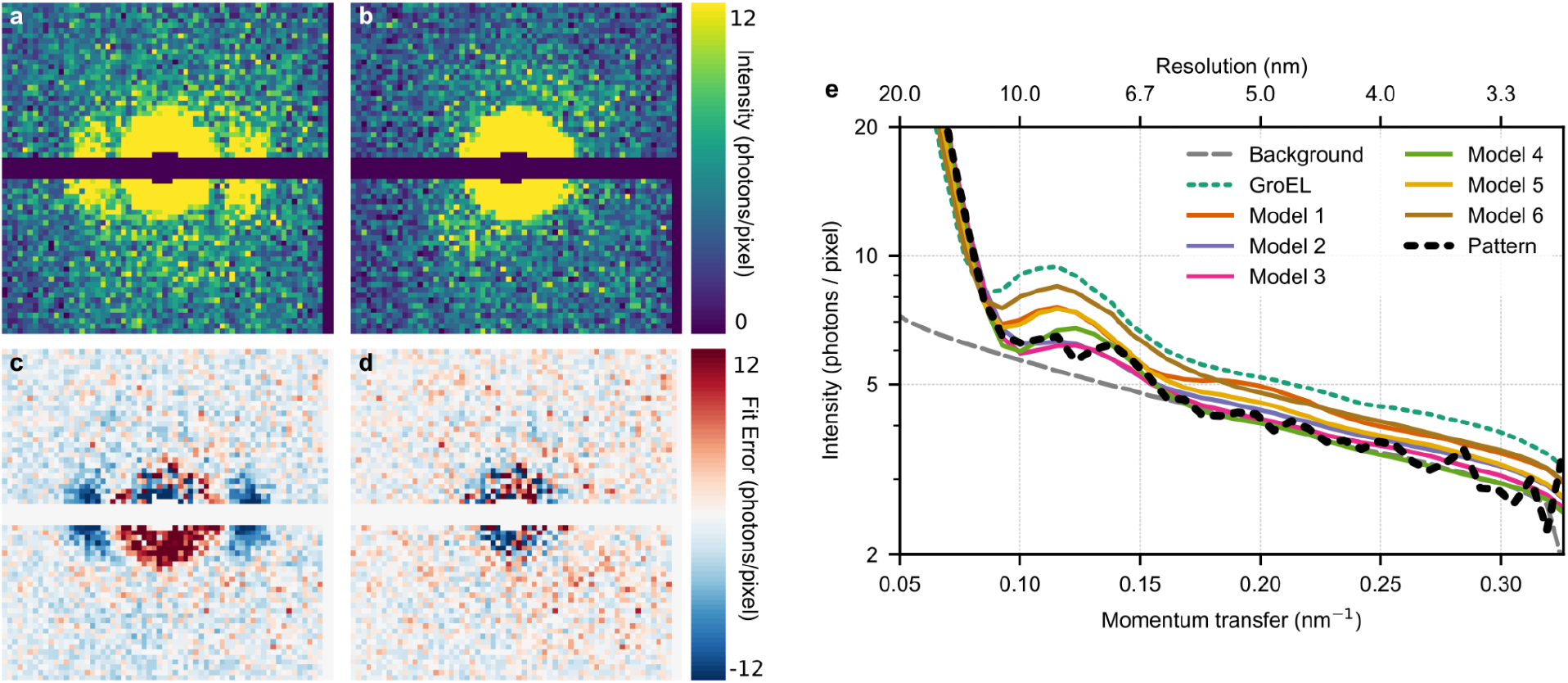
Pattern comparisons. **a**, Simulated diffraction from the dry GroEL particle in the orientation that best matches the measured pattern. Patterns are plotted with Poisson noise. **b**, A significantly better fit is achieved from density model 3. **c** and **d**, Fit error between the measured pattern compared to **c** the simulated diffraction from the dry GroEL particle and **d** model 3. **e**, Radial average of the background, the best fitting diffraction from the crystal structure of GroEL (dotted green line), each density model (solid coloured lines) and the measured pattern (dotted black line). The dry molecule predicts too much intensity outside of the central speckle whereas water models 2, 3 and 4 follow the data much closer. The images (**a**, **b**, **c** and **d**) and the pixels of **e** correspond to 64×64 downsampled patterns.

The radial average of the pattern and the best fit for the different density models showed a better fit for all new models compared to the original structure, with models 2, 3 and 4 giving the best results (**Fig. 4c**). The total residual error between the simulation and the experimental pattern also confirmed that model 3 was the best fit with an error of 197 photons. The simulation from model 3 is shown in **Fig. 4b** and the oriented model is shown in **Fig. S7**. The signal in the radial average of the pattern drops down to the background level at a resolution of about 6 nm.

Not only are these results consistent with diffraction from a GroEL molecule, which is the first example of interpretable X-ray diffraction being collected from a single protein, but they also suggest that the aerosolized GroEL particle contained an extra density in the otherwise hollow centre at the time of interaction.

The overall size and shape of our sample match that of the crystal structure quite well, unlike earlier studies using a combination of ion mobility analysis and mass spectrometry^31^ which have observed an unusually high compaction of GroEL in the gas phases. The difference is likely due to the different experimental conditions. In our case GroEL was quickly neutralised after electrospray and not actively dried, while the compaction was seen for dry particles with charges up to z=70, which is likely to affect the structure. This suggests that hydration and charge state are important to preserve the GroEL structure in FXI experiments.

From our modelling, we also concluded that of the 30,500 photons in the pattern, only 13,800 originated from the sample and 16,700 originated from the background scattering. This highlights the importance of continued efforts to further reduce background scattering from the injection gas in such experiments.

The pattern fittings showed that the photon fluence at the sample was 280 μJ/μm^2^. This aligns well with the maximum fluence expected from the pulse given a measured pulse energy of 6.6 mJ before the focusing optics and the focus profile and transmission of the beamline (see Methods). It suggests that this particular GroEL molecule interacted with a region of the pulse that was almost at the peak.

When the first XFELs were constructed, one of the main promises was the prospect of diffraction studies of single proteins using the so-called “diffraction before destruction method” that could take advantage of the ultrafast time-resolution enabled by this new generation of light sources. However, concerns were raised on whether the proteins’ structure would survive the transition to the gas phase and, even if it did, whether the signal would be strong enough to be visible above the background noise. In this paper, we have been able to address these concerns by reporting the first X-ray diffraction pattern collected from a single protein.

The signal in this pattern is weak, but the distinct geometry of the GroEL complex is distinguishable above the background noise. Furthermore, the signal matches well with the predicted signal from a model of GroEL with extra density added to the central cavity. At this resolution, it cannot be determined if the extra density is made up of water or something else.

Simulations have shown that residual water molecules are vital for the stability of proteins in the gas phase^32^. A significant amount of water attached to GroEL in our experiment would, without doubt, contribute to keeping its structure preserved during the transition to the gas phase. The presence of water around the sample is also predicted to delay radiation damage to the sample by acting as a sacrificial tamper^33^. Large amounts of solvent might introduce problems for 3D orientation recovery and subsequent merge of a large dataset. These problems will however be limited to the same resolution as the size of the fluctuations in solvent distribution between the samples, which for water is expected to be small^34^.

The factors that currently prevent FXI from determining full 3D structures are the low signal-to-noise ratio due to the strong background and the low data rate. Since most of the background originated from the injection gas, we identify this as a major target for future development. Potentially, better shielding of the gas and a transition to a low-Z alternative such as helium could improve the signal-to-background ratio by more than tenfold. The availability of a 4.5 MHz DSSC imaging detector of megapixel size^35^ at the SQS instrument will allow us to exploit the 4.5 MHz pulse repetition frequency within one pulse train of the XFEL, yielding multiple opportunities for a hit in each pulse train. Furthermore, the vetoing capability^36,37^ of the DSSC detector has the potential to improve the fraction of interpretable diffraction images from a few percent to around 30 percent when EuXFEL is running at its full capacity of 27 kHz.

Here we have presented the first interpretable X-ray diffraction pattern from a single protein, frozen in time by the femtosecond X-ray pulse, and experimentally demonstrated that the concept of diffraction before destruction extends to single proteins. This single pattern represents an important step towards solving 3D protein structures with the method of diffraction before destruction and shows that several of the hurdles can indeed be overcome. With higher data rates, many such patterns can map out the structure and function of dynamic proteins with the staggering time-resolution enabled by XFELs.

## Methods

### Beamline and instrument setup

The EuXFEL was tuned to a photon energy of 1200 eV corresponding to a wavelength of 1.03 nm. The focus size was estimated to be 2 μm × 2 μm based on wavefront sensor measurements (**Fig. S8**). The total energy of each X-ray pulse was measured before any beamline optical element using one of the X-ray gas detectors available at the beamline^38^ and found to hover around 6.5 mJ. Using the wavefront sensor measurements (**Fig. S8**) we estimated the fluence at the interaction region. We assumed that the field of view of the sensor captures the vast majority of the photons present in the beam. Using the measured pulse energy and a beamline transmission of 46% (measured subsequently), we estimated the maximum fluence across the sensor for each of the five different wavefront measurements. The average of those estimates was 232±62 μJ/μm^2^. The XFEL was run at one pulse per train giving a repetition rate of 10 Hz.

### Sample injection

Individual proteins were transferred into the gas phase and transported into the X-ray interaction region as described in Bielecki *et al.^24^*. The sample solution consisted of GroEL proteins with a concentration of about 150 nM in an ammonium acetate buffer. Nebulization of the protein solution took place with an electrospray nozzle which produces initial droplets with diameters between 80-400 nm depending on the sample flow rate. The charged droplets emanating from the electrospray nozzle were neutralised by an X-ray source (Hamamatsu L12645) that ionised the sheath gas transporting the droplets.

The electrospray capillary had an inner diameter of 40 μm, an outer diameter of 360 μm, and the sample flow rate was adjusted by controlling the overpressure in the sample compartment with a remotely controllable differential pressure regulator (Bronkhorst P-506C-4K0D-TGD-33-V delta P pressure gauge controlling an F-001AI-IIU-33-V regulating valve). The tip of the capillary had been ground to a 30-degree cone with a final tip diameter of 100 μm. The droplet diameter could be controlled from 80 nm at 0.25 psi overpressure to 400 nm at 10 psi overpressure.

Monodispersity and size of the sample after nebulisation were both monitored before, and during the measurements, with an SMPS (TSI SMPS 3938) consisting of a DMA coupled to a condensation particle counter. To minimise the salt layer on the sample surface, while still maintaining a stable Taylor cone, an overpressure of 1 psi had to be applied to the sample reservoir used, resulting in initial droplets with a diameter of approximately 110 nm.

The neutralised droplets were transported into the X-ray interaction region through an aerodynamic lens, creating a particle beam as described in Hantke *et al.^25^*. Excessive gas flow from running the electrospray was removed in two skimmer stages. As a result, the 1 bar pressure at the electrospray was reduced to 30 mbar after the first skimmer, and the entrance pressure to the aerodynamic lens was 0.6 mbar after the second skimmer stage.

The beam of injected particles was intercepted by the pulse train of the XFEL. To optimise the position of the particle beam, a sucrose solution was injected, creating tiny sucrose spheres, and the hit-rate on the spheres was used as a feedback parameter.

### Detector and Data Processing

Diffraction data were collected with the EuXFEL pnCCD detector^27^ running in high-gain mode. This setup allows for a maximum full-period resolution of 4 nm determined by the scattering-angle at the edge of the detector. Since the detector cannot keep up with the pulse frequency within the pulse trains, we were limited to the 10 Hz frequency of the pulse trains themselves. Each pnCCD sensor panel is made up of a grid of 512 × 1024 pixels each with a size of 75 μm × 75 μm. The two panels were both placed 15 cm downstream of the interaction region and with a gap of 3.7 mm to allow the direct beam to pass through. The exact translation of the detector panels was optimised using strongly diffracting sucrose particles and the understanding that this diffraction adheres to Friedel symmetry.

Pedestal data were collected regularly throughout the experiment when the beam was off and were subtracted from each readout. In addition, a common mode correction was applied to each line of each detection plane for each individual image. This correction is performed by subtracting the median of the pixel values in each line from all values of the same line and is possible when the photon density is low, like in our case.

The slope of the relation between photon energy and ADU of each detector pixel, called the gain, varies slightly from line to line since each line has its own amplifier. In addition, along a line, the measured energy might decline due to the charge transfer inefficiency. To handle both these effects we determined a unique gain for each pixel. This value was found by constructing a histogram of the signal detected in all images in a particular pixel (**Fig. S6**) and subsequently fitting a Gaussian function to the peak corresponding to zero photons and subsequently fitting another Gaussian function to the much smaller peak corresponding to a single photon. The distance between the peaks must then correspond to the photon energy of 1200 eV.

The detector signal provided in units of ADU was converted into photon counts by rounding each pixel readout to its closest integer. To filter out the contribution from fluorescence in the range from 200 eV to 600 eV readout values up to 900 eV were rounded down to zero instead of up.

For each readout, the number of lit pixels was calculated as the number of pixels with a non-zero photon count. Hits were identified as any readout where the number of lit pixels was larger than 16.

The average background was estimated from 32,000 readouts (**Fig. S9**) where the injector was running but without any sample, thus including the contribution of the scattering from the gas used for injection.

Before analysis, each diffraction pattern was downsampled to a size of 128 × 128 pixels. The downsampling was done after the conversion to discrete photons since the combined readout noise in one superpixel would otherwise be much larger than the photon energy. Additional downsampling to a final size of 64 × 64 was performed before plotting to make the features of the diffraction patterns more clear.

Water models were generated by solvating the GroEL structure (PDB entry 1SS8^30^) using the gmx solvate function in GROMACS^39^. Water molecules were removed if they fell outside of a cylinder of varying size. The top and bottom of the cylinders were also pruned to match the shape of the protein. The code for generating these models and the PDB files for them are made available (see Code Availability).

Template diffraction patterns were simulated with Condor^40^ using the wavelength and detector geometry from the experiment. The output without Poisson noise was used in the further analysis. The protein orientations were distributed evenly in rotation space by choosing quaternions that evenly sample the cells of the 600-cell similarly to Loh *et al.^41^*. Each edge in the 600-cell was subdivided 8 times which yields 25,680 different orientations and corresponds to an angle of 6.8 degrees between adjacent orientations.

For the template matching, each template was combined with the average background with a variable scaling term for the fluence of the signal and background respectively. These scaling terms were used as fitting parameters in a least-square optimization implemented in the scipy function leastsq^42^. The goodness of fit was then compared between all templates to identify the best orientation.

The residual error, *E*, or goodness of fit, is defined as

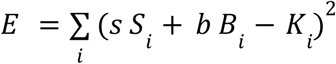

where *i* is the pixel index and *S* is the simulated template, *B* is the average measured background and *K* is the measured pattern. The parameters *s* and *b* are the fitting parameters and describe respectively the intensity of the pulse at the sample and total intensity of the pulse.

### Sample purification

Lyophilized *E. coli* GroEL (C7688) was purchased from Sigma–Aldrich (Solna, Sweden), purified and prepared for electrospray injection as described in Freeke *et al.^43^*, but with no acetone precipitation step and with one step of size exclusion chromatography.

### Characterization of GroEL samples by DMA

The stability of GroEL against dissociation was determined using DMA combined with the same electrospray conditions as the particle injection for the main experiment. Here, a narrow peak at 16 nm was recorded which suggests that GroEL is stable under the XFEL injection conditions. A second larger peak was also detected at a smaller diameter that corresponds to contaminants from empty droplets aggregating to a ball (**Fig. S3**).

### Characterization of GroEL samples by cryo-EM

For cryo-EM, vitrified grids were prepared by applying 4 μl of the GroEL sample onto glow-discharged, 200 mesh R2/2 Quantifoil grids, blotted for 4 seconds at blotforce 4. Grids were plunge-frozen into a 37:63 (v/v) mixture of ethane/propane cooled to liquid nitrogen temperature using a Vitrobot Mark IV instrument (ThermoFisher Scientific) at 95% humidity and 4 °C. Samples were imaged at a nominal magnification of x120,000 using a Talos Arctica (ThermoFisher Scientific) transmission electron microscope operating at 200 kV accelerating voltage from a field emission gun (X-FEG) source. Movies were recorded on a Falcon 3EC electron counting direct detector (ThermoFisher Scientific) yielding a final pixel size of 0.96 Å^2^ on the specimen level. A total of 497 movies were collected in dose-fractionation mode using EPU software (ThermoFisher Scientific) with a total dose of 40 e^−^/Å^2^ for each micrograph, and 1e^−^/Å^2^/frame.

#### Cryo-EM data processing

Image processing was done in a combination of RELION 3.1^44^ and cryoSPARC^45^. Movies were processed using MotionCorr 2^46^ as implemented in RELION 3.1 for motion correction and gCTF^47^ for CTF correction.

#### Cryo-EM data analysis: sample composition analysis

Laplacian picking in RELION 3.1 considers the fact that for a quality assessment a bias-free, reference-free particle picking is needed. For this both threshold and particle size were optimised until nearly all particles, visible by eye, were picked up by the program, and as little as possible noise was included, although some error was still present (see **Fig. S2** for an example). This resulted in a total of 47,154 particles picked with a threshold of 2 and a picked particle size between 120 and 900 Å. These particles were subsequently classified in 200 classes in cryoSPARC^45^.

Only classes containing GroEL particles were submitted to heterogeneous refinement in cryoSPARC. For this, two references were supplied, one for the dual- and one for the single-ring complex. The first was an intermediate low-resolution map that was constructed during this project (see next section), aligned to D7 symmetry. The second was created based on a single-ring from the PDB structure 5W0S^48^ by using the molmap function in Chimera 1.15^49^ with a resolution of 20 Å. This map was subsequently resampled to the correct box and pixel size in Chimera 1.15, followed by alignment in RELION 3.1 to C7 symmetry (to centre and prepare for symmetry application). Following heterogeneous refinement, the two groups of particles were submitted to another round of 2D classification, to make sure that the separation had been thorough (see **Fig. S3**). No classes belonging to the other complex were detected, but a few classes containing noise and smaller pieces of the complex were removed prior to calculating the ratio between single- and dual-ring particles in the sample. A selection of top views from the 2D classes of the dual-ring group of particles was used for **Fig. S4**.

Those classes containing small proteins were 2D cleaned and the more prominent classes were subjected to initial 3D model generation in cryoSPARC. Ten low-quality 3D models were generated and they were all of similar size. Since this size was comparable to monomeric GroEL, a 3D refinement in cryoSPARC and a 3D classification in RELION 3.1 was performed. The reference was created based on a monomer from the PDB structure 5W0S by using the molmap function in Chimera 1.15 with a resolution of 20 Å. This map was subsequently resampled to the correct box and pixel size in Chimera 1.15. Neither analysis yielded a map with improved density. As the identity of these small particles is not relevant to the XFEL experiments they were not further analysed.

#### Cryo-EM data analysis: high-resolution model

A deep-learning-based picking in crYOLO^50^ to allow for precise picking of intact GroEL particles, resulted in a total of 14,232 particles that were imported into RELION 3.1. These were subjected to 2D classification into 50 classes and the best 10 classes were used for 3D classification into four classes with D7 symmetry in RELION 3.1. The best class included 1929 particles corresponding to the dual-ring complex and was refined with D7 symmetry and postprocessing leading to a final map resolved to 4.6 Å as shown in **Fig. S5**.

## Code Availability

Simulations were performed with the open-source software package Condor^40^. Software to perform the template matching and all auxiliary software for gain correction, and hit finding are available at https://github.com/ekeberg/Ekeberg2022GroEL.

## Data availability

A total of 94750 detector images were deposited on the Coherent X-ray Imaging Data Bank (CXIDB)^51^ under [ID 187](https://cxidb.org/id-187.html). This includes sample runs (83600 images), detector calibration runs (3750 images), runs with only the X-ray beam (1200 images) and X-ray beam, sample delivery gas but without sample (6200 images). The DOI for the original data at the EuXFEL is https://doi.org/10.22003/XFEL.EU-DATA-002146-00.

## Acknowledgements

We acknowledge European XFEL in Schenefeld, Germany, for provision of X-ray free-electron laser beam time at the SQS instrument and would like to thank the staff for their assistance. We acknowledge the use of the XBI biological sample preparation laboratory, enabled by the XBI User Consortium. We acknowledge valuable discussions with Erik G. Marklund. The results of the work were obtained using Maxwell computational resources operated at Deutsches Elektronen-Synchrotron (DESY), Hamburg, Germany. Part of this work was performed at the Multi-User CryoEM Facility at the Centre for Structural Systems Biology, Hamburg, supported by the Universität Hamburg and DFG grant numbers (INST 152/772-1|152/774-1| 152/775-1|152/776-1|152/777-1 FUGG). We acknowledge the support of funding from: Cluster of Excellence ‘CUI: Advanced Imaging of Matter’ of the Deutsche Forschungsgemeinschaft (DFG) - EXC 2056 - project ID 390715994; ERC-2013-CoG COMOTION 614507; NFR 240770; Fellowship from the Joachim Herz Stiftung (P.L.X.); P.L.X. and H.N.C. acknowledge support from the Human Frontiers Science Program (RGP0010/2017); J.H. acknowledges support from the European Development Fund: Structural dynamics of biomolecular systems (ELIBIO) (CZ.02.1.01/0.0/0.0/15_003/ 0000447) EMBO long-term fellowship (ALTF 356-2018) awarded to L.E.F.; the Röntgen-Ångström Cluster (2015-06107 and 2019-06092); the Swedish Research Council (2017-05336, 2018-00234 and 2019-03935); the Swedish Foundation for Strategic Research (ITM17-0455).

## Author contributions

T.E, K.A., H.N.C., Lars.G., J.H, J.K., D.W. and F.R.N.C.M. conceived and designed the experiment. L.A.E., L.H.G., D.H., A.K.S. and P.L.X. prepared and characterised the sample. L.E.F, W.L. and C.S. performed cryo-EM measurements and analysis. J.B., J.K., O.K, J.L., A.P.M., A.K.S. and L.W. developed and operated the sample delivery equipment. D.A., K.A., B.J.D., T.E., D.E.G., Luca.G., A.I., J.K., R.K., F.R.N.C.M., C.N., E.S., J.A.S., N.T., I.A.V., and T.W. contributed to software development, data processing and analysis. R.B., M.M., T.M., Y.O., D.R., P.S. and S.U. designed and operated the SQS instrument at EuXFEL. R.H., H.Y. and M.K integrated and operated the pnCCD detector. The manuscript was written by T.E. and F.R.N.C.M. with input from all authors.

