## Supplementary Information for "Observation of a single protein by ultrafast X-ray diffraction"

Tomas Ekeberg et al.

### Supplementary Information

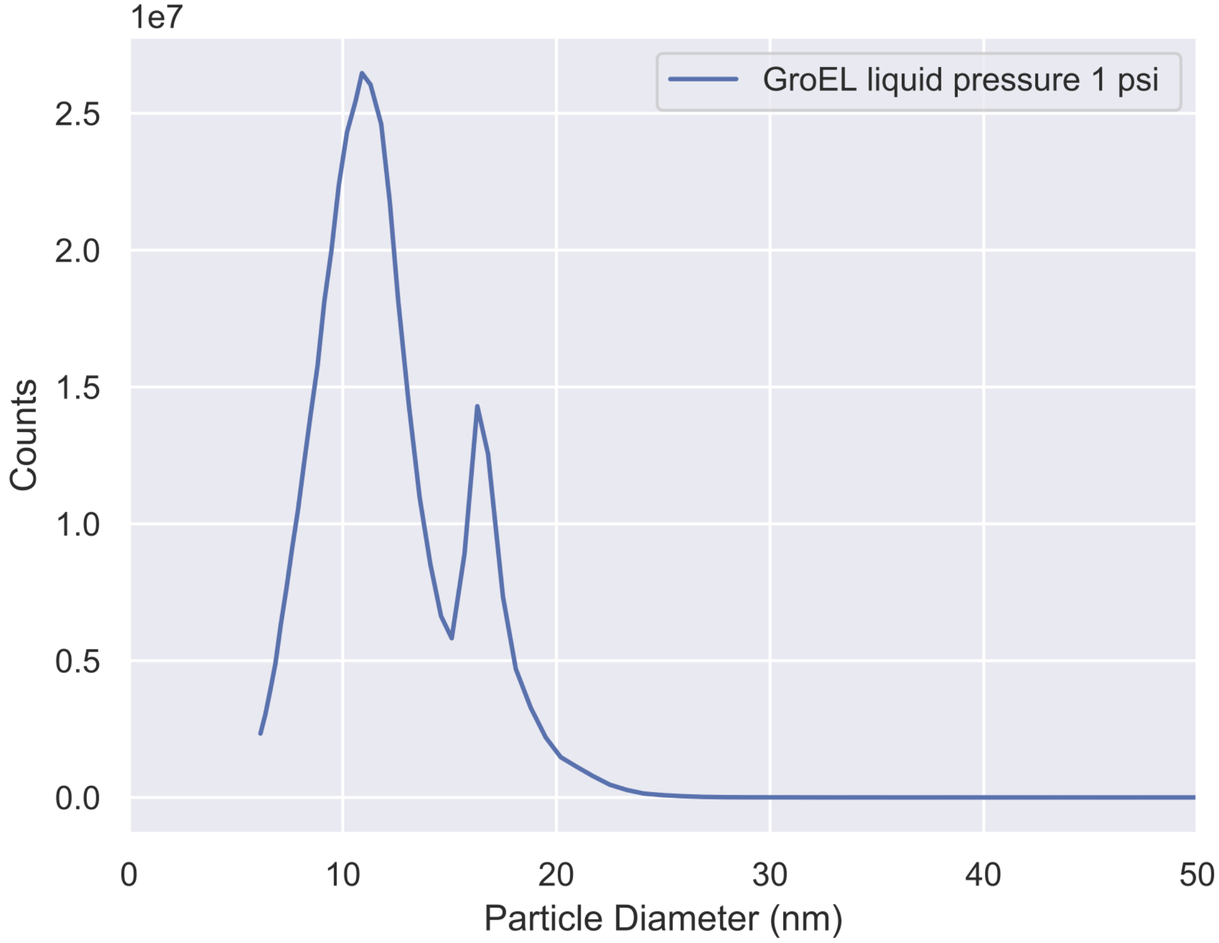


**Figure S1 | Differential mobility analyzer(DMA) measurement of the sample.** Spectrum of the particle diameters measured by the DMA, when running the electrospray in the same conditions that gave rise to the patterns described. The peak around 11 nm corresponds to the impurities in droplets without GroEL while the peak around 16 nm corresponds to a droplet with a GroEL complex.

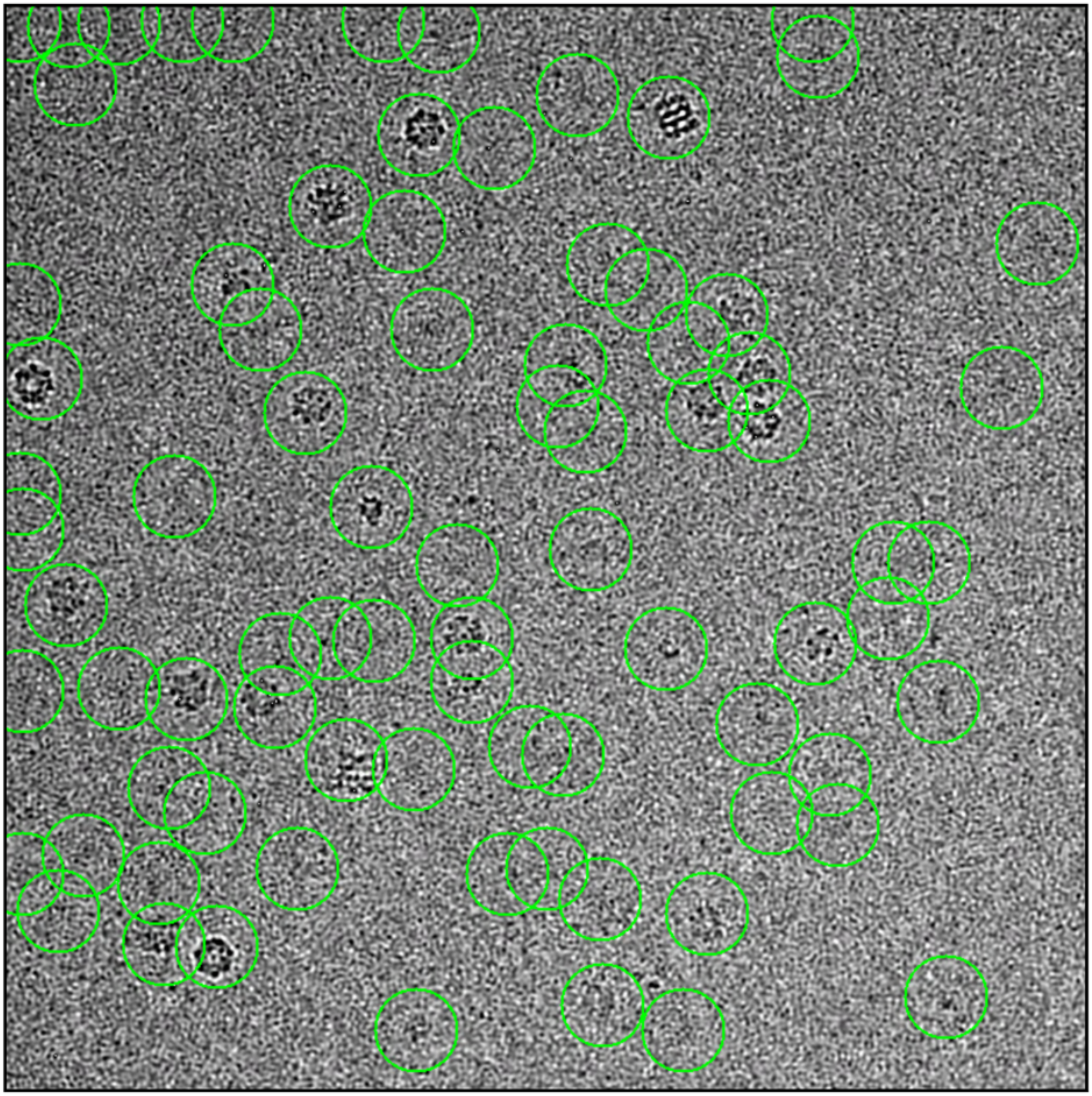

**Figure S2 | Particle Selection based on the Laplacian Picking algorithm is shown in an example micrograph.** Reference free particle selection with the Laplacian Picking algorithm from Relion 3.1 picks all intensities between 12 and 90 nm. The green circle is 300 Å in diameter. As can be seen in this example micrograph, most particles are picked as well as several small intensities in the background which can be noise or small proteins. This demonstrates unbiased picking, ideal for sample quality analysis.

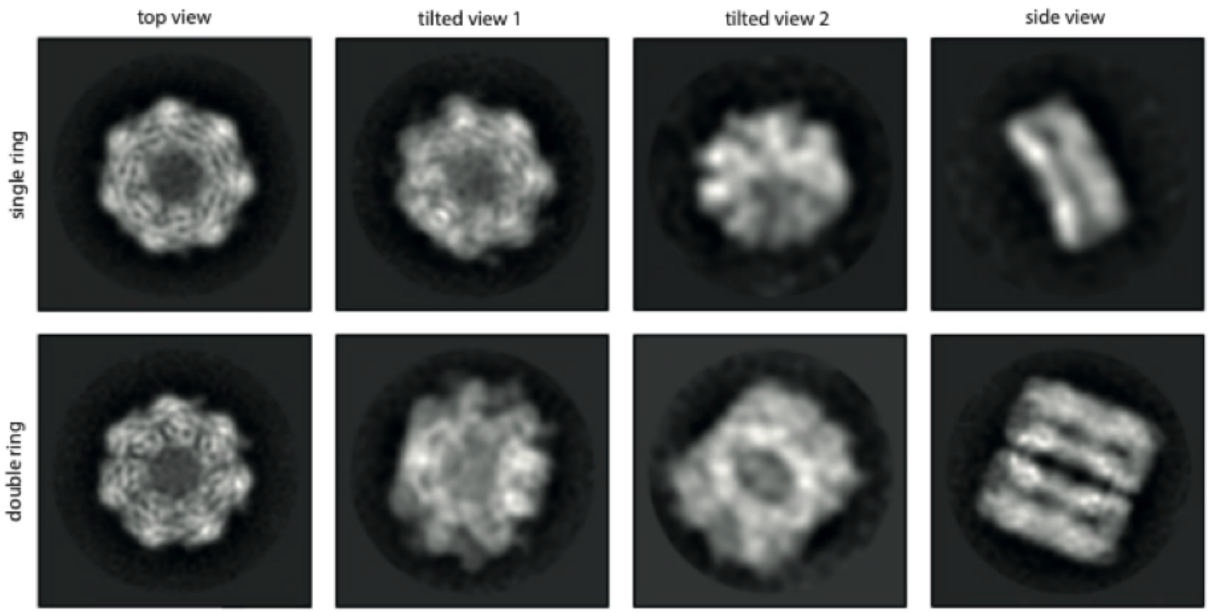


**Figure S3 | Representative 2D class averages from Cryo-EM images of the sample.** Sample quality analysis shows the presence of single- and dual-ring GroEL complexes. Eight representative 2D class averages from Cryo-EM images of the sample are shown. From projection images, it cannot be concluded whether these two different top views correspond to single-ring or dual-ring complexes. 2D cleaning of the picked dataset (**Fig. S2**) removed most of the particles that contained small protein or noise, and subsequent 3D heterogeneous refinement against a single- and a dual-ring reference, followed by further 2D cleaning allowed us to reduce the dataset to 3454 particles, of which 869 particles (32%) and 2676 particles (68%) were assigned to single- or dual-ring complexes respectively.

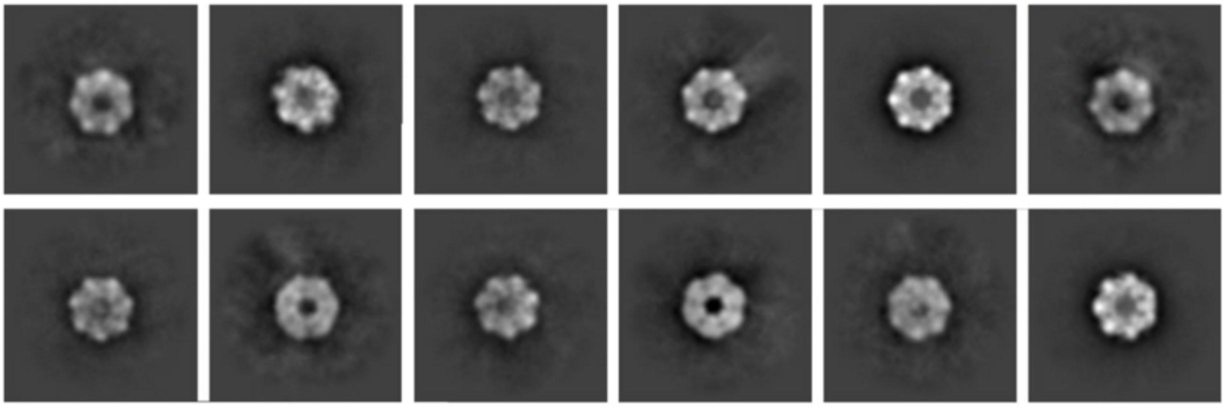

**Figure S4 | Gallery of top view 2D classes corresponding to dual-ring GroEL.** Gallery of top view 2D classes corresponding to dual-ring GroEL. Images were selected from the last round of 2D cleaning after heterogeneous refinement from the subset of particles corresponding to the GroEL dual-ring. Density can be observed in the centre of several classes, likely from attached proteins.

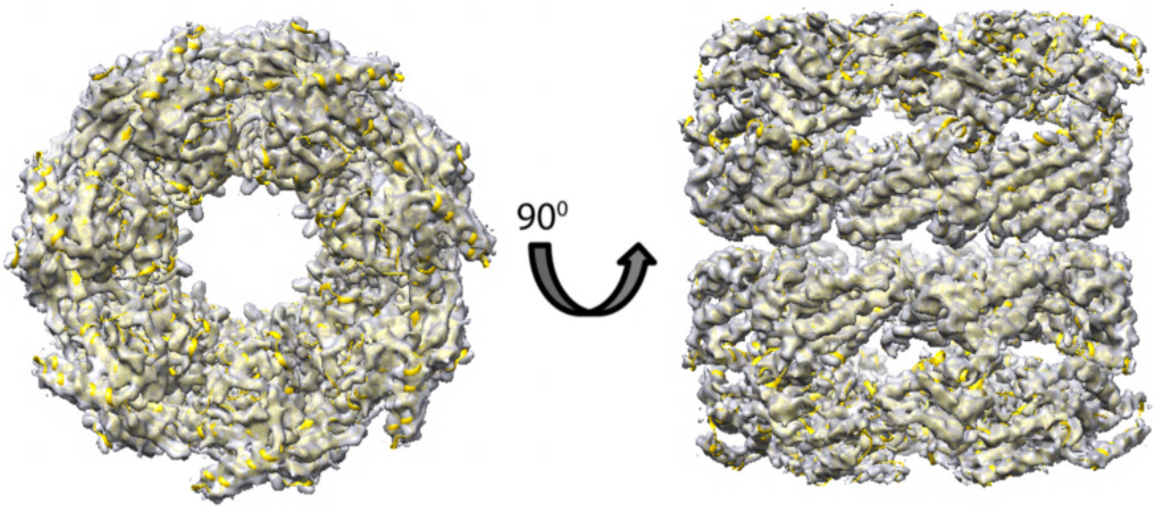


**Figure S5 | Final 3D map at 4.6 Å resolution fitted to the PDB structure (5W0S).** Final 3D map of GroEL dual-ring protein at 4.6 Å resolution with PDB structure (5W0S) fitted inside. No apparent differences were observed between map and model. As D7 symmetry was enforced during the 3D reconstruction, details that do not comply with the symmetry group, including any density within the rings, were averaged out and became invisible in the final 3D density.

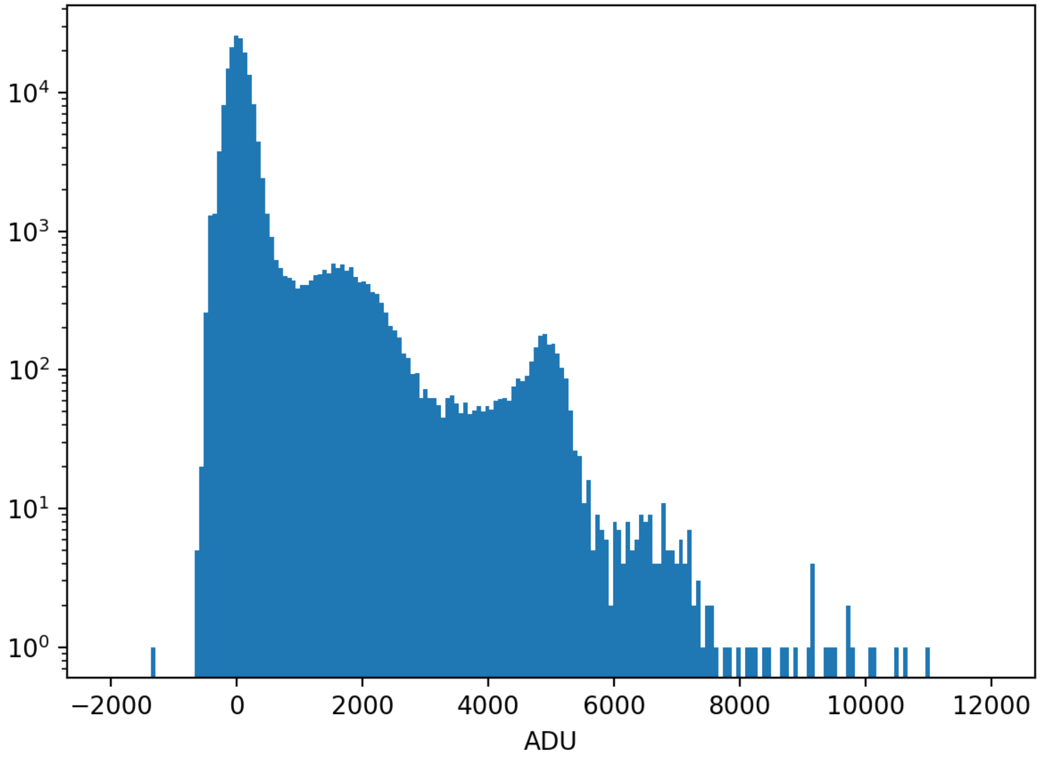


**Figure S6 | Histogram of the detector signal for a single pixel.** The sharp peak around 5000 ADU corresponds to a single X-ray photon at 1200 eV. The broad peak around 2000 ADU corresponds to the fluorescence signals from the $K_{\alpha 1}$-shells of carbon (277 eV), nitrogen (392 eV) and oxygen (525 eV).

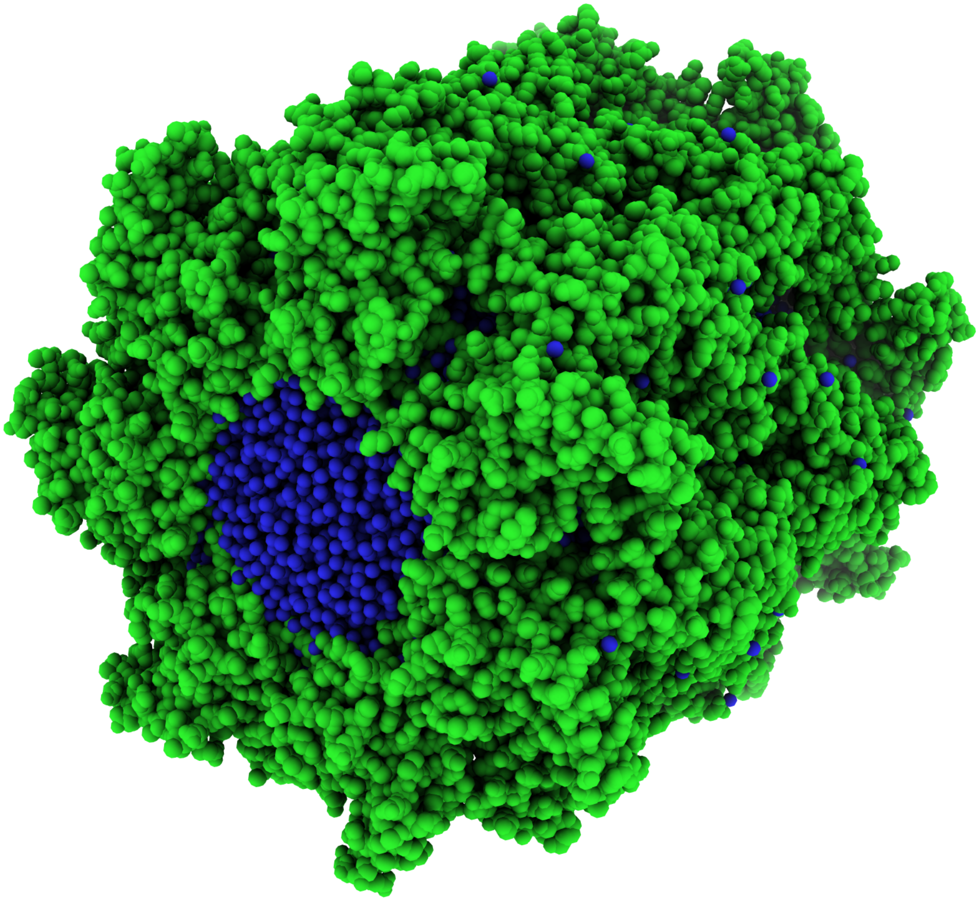

**Figure S7 | Orientation of the GroEL in the X-ray beam.** Model of GroEL in the orientation that best matches the experimental diffraction, corresponding to the orientation of the density model 3 that gives rise to the diffraction pattern in **Fig. 4b**. The X-ray beam direction is defined to be directly into the page.

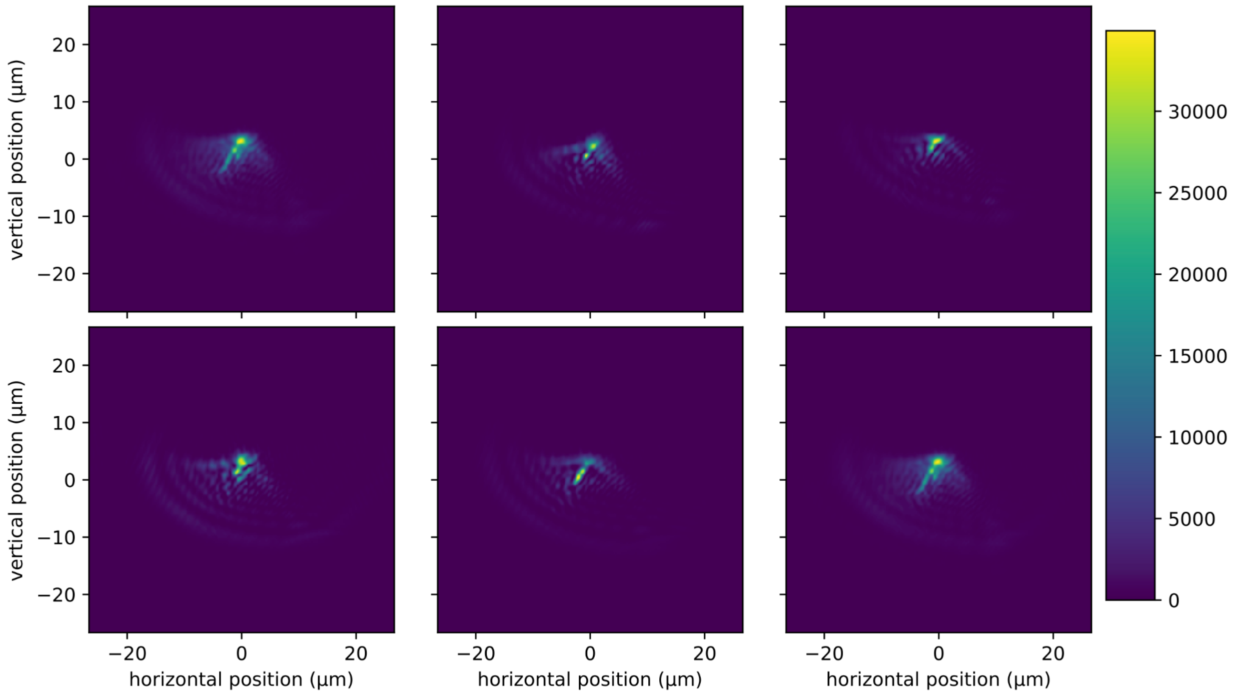


**Figure S8 | Focal plane intensity distribution of different shots.** Single-shot intensity distribution, of the focal plane, of five different shots. The bottom right corresponds to the average of the five shots. There was a variation in the focal plane intensity distribution from shot to shot, with an associated change in the focal spot. From the average intensity, the full width at half maximum of the focus was estimated at 1.7 µm x 2.3 µm.

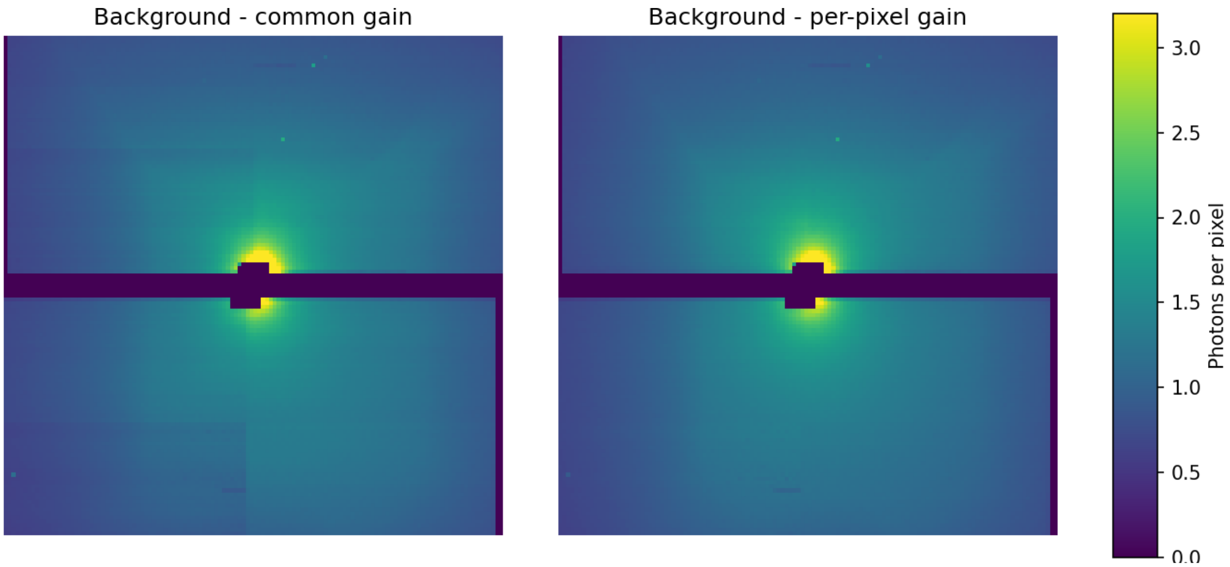


**Figure S9 | Average photon background with and without per-pixel gain.** When assuming the same gain for all the pixels, the boundaries between different ASICs in the detector are visible, as shown on the left-hand side. By calibrating the gain of each pixel, using pixel-level histograms, one obtains a smooth image with no visible boundaries, as shown in the right, which matches the expectation of what the gas background should look like.
